# Lateral diffusion of NKCC1 contributes to neuronal chloride homeostasis and is rapidly regulated by the WNK signaling pathway

**DOI:** 10.1101/2022.12.11.519958

**Authors:** Etienne Côme, Simon Blachier, Juliette Gouhier, Marion Russeau, Sabine Lévi

**Affiliations:** INSERM UMR-S 1270, Sorbonne Université, Institut du Fer à Moulin, 75005, Paris, France

**Keywords:** Hippocampal neurons, chloride homeostasis, GABAergic transmission, lateral diffusion, clustering, protein trafficking, signaling, single molecule localization microscopy

## Abstract

An upregulation of the Na^+^-K^+^-2Cl^-^ co-transporter NKCC1, the main chloride importer in mature neurons, can lead to depolarizing/excitatory responses mediated by GABA_A_ receptors and thus to hyperactivity. Understanding the regulatory mechanisms of NKCC1 would help prevent intra-neuronal chloride accumulation that occurs in pathologies with defective inhibition. The cellular and molecular regulatory mechanisms of NKCC1 are poorly understood. Here, we report in mature hippocampal neurons that GABAergic activity controls the membrane diffusion and clustering of NKCC1 via the chloride-sensitive WNK1 kinase and the downstream SPAK kinase that directly phosphorylates NKCC1 on key threonine residues. At rest, this signaling pathway has little effect on intracellular Cl^-^ concentration but it participates to the elevation of intraneuronal Cl^-^ concentration in hyperactivity condition associated with an up-regulation of NKCC1. The fact that the chloride exporter KCC2 is also regulated in mature neurons by the WNK1 pathway indicates that this pathway will be a target of choice in the pathology.

## Introduction

Upon activation by GABA, GABA type A receptors (GABA_A_Rs) open a selective chloride/bicarbonate conductance. The direction of chloride (Cl^-^) flux through the channel depends on transmembrane Cl^-^ gradients. Therefore, Cl^-^ homeostasis critically determines the polarity and efficacy of GABAergic transmission in the brain. Pharmaco-resistant epilepsies are often associated with altered Cl^-^ homeostasis [1]. It is therefore crucial to discover novel mechanisms regulating neuronal Cl^-^ homeostasis that may help develop new and efficient treatment for these forms of epilepsy and other diseases associated with impaired inhibition, such as neuropathies and neuropsychiatric disorders [2].

The increase in [Cl^−^]_i_ and subsequent depolarized shift in the reversal potential of GABA (E_GABA_) observed in adult epilepsy models are often attributed to reduced expression/function of the neuronal K^+^-Cl^-^ cotransporter KCC2, responsible for Cl^-^ export [3]. Furthermore, up-regulation of the Na^+^-K^+^-Cl^-^ cotransporter NKCC1, which imports chloride into neurons, also contributes to the increase in [Cl^-^]_i_ and depolarization of E_GABA._ This has been observed in the subiculum of patients with temporal lobe epilepsy (TLE) [4-5] as well as in several *in vivo* and *in vitro* rodent models of epilepsy such as the glioma-induced [6], the traumatic brain injury-induced [7] models of epilepsy, and the kainic acid and pilocarpine epilepsy models in tissue slices [8-10]. Targeting NKCC1 in epilepsy with the inhibitor bumetanide is protective: it abolished glioma-induced seizures in rats [6], it reduced the frequency of interictal-like activities [4], the duration of ictal activities [10], and the sprouting of mossy fibers [9].

Key cellular and molecular mechanisms regulate the membrane turnover of KCC2. They involve activity-dependent regulation of the transporter membrane turnover and endocytosis [11-12]. KCC2 is rapidly down-regulated by enhanced neuronal activity and glutamatergic neurotransmission in mature neurons [13-15]. NMDA receptor-induced Ca^2+^-influx leads to protein phosphatase 1 (PP1)-dependent KCC2 Serine S940 (S940) dephosphorylation and C-terminal domain cleavage by Ca^2+^-activated protease calpain [14, 16-17]. This in turn leads to increased lateral diffusion, endocytosis and degradation of KCC2 [13-14]. GABAergic signaling also tunes KCC2 at the neuronal membrane through GABA_A_R activity and Cl^-^-dependent phosphorylation of KCC2 Threonine 906 and 1007 (T906/1007) residues [18]. Cl^-^ acts as a second messenger in this regulation by tuning the activity of the Cl^-^ sensing With No lysine (K) serine-threonine kinase WNK1 and its downstream effectors Ste20 Proline Asparagine Rich Kinase (SPAK) and Oxydative Stress Response kinase 1 (OSR1) [19].

Interestingly, this signaling not only promote KCC2 T906/T1007 but also NKCC1 T203/T207/T212 phosphorylation [20]. This results in a powerful modulation of Cl^-^ transport by inhibiting KCC2 and activating NKCC1; both regulations leading to elevation in intracellular Cl^-^ level in non-neuronal cells [21]. Therefore, inhibiting KCC2 and NKCC1 phosphorylation mediated by the WNK signaling might normalize the membrane expression/function of transporters, thereby preventing the intracellular Cl^-^ buildup in epilepsy and the intensity or emergence of epileptic seizures.

Our team has shown the contribution of lateral diffusion in the rapid control of KCC2 membrane stability and Cl^-^ neuronal homeostasis in response to changes in neuronal activity [13,18]. The transporter alternates between periods of confinement within clusters near synapses and periods of free movement outside the clusters. The free transporter can then be targeted to the endocytic wells where it is internalized and then degraded or recycled to the plasma membrane. These two pools of transporters are in dynamic equilibrium, allowing the fine-tuning of synapses in response to local fluctuations in synaptic activity [11-12]. Since changes in KCC2 mobility occur within tens of seconds [18], lateral diffusion is probably the primary cellular mechanism modulating KCC2 membrane stability.

Recently, we showed with confocal microscopy that NKCC1 is targeted to the axonal and somato-dendritic plasma membrane and that it forms clusters at the periphery of synapses of hippocampal neurons [12]. Quantum Dot-based Single Particle Tracking (QD-SPT) in living cultured hippocampal neurons indicated that NKCC1 explore large areas of the somato-dendritic extrasynaptic membrane while others are confined near excitatory and inhibitory synapses [12]. Transitions between NKCC1 confinement at/near synapses and less constrained diffusion in extrasynaptic areas is reminiscent of KCC2 diffusion [13,18]. Therefore, similar to KCC2, NKCC1 responds to the “diffusion-trap” mechanism [12]. A rapid activity-dependent regulation of NKCC1 diffusion/clustering, may thus locally affect its function, and in turn GABA signaling.

Here we show that neuronal GABAergic activity rapidly regulates lateral diffusion and membrane clustering of NKCC1 in mature hippocampal neurons in culture. Blocking or activating GABA_A_ receptors with muscimol or gabazine confines the transporters to the dendritic membrane. GABAergic activity would control diffusion and clustering of NKCC1 in the dendrite via the chloride-sensitive kinase WNK1 and the downstream kinases SPAK and OSR1 that would directly phosphorylate NKCC1 at key threonine phosphorylation sites T203/T207/T212. Our results indicate that this regulation would be particularly effective in regulating NKCC1 and chloride homeostasis in the somato-dendrtic compartment under hyperactive conditions, highlighting the value of targeting the WNK1/SPAK/OSR1 pathway in inhibition-defective pathologies.

## Materials and Methods

For all experiments performed on primary cultures of hippocampal neurons, animal procedures were carried out according to the European Community Council directive of 24 November 1986 (86/609/EEC), the guidelines of the French Ministry of Agriculture and the Direction Départementale de la Protection des Populations de Paris (Institut du Fer à Moulin, Animalerie des Rongeurs, license C 72-05-22). All efforts were made to minimize animal suffering and to reduce the number of animals used. Timed pregnant Sprague-Dawley rats were supplied by Janvier Lab and embryos were used at embryonic day 18 or 19 as described below.

### Neuronal culture

Primary cultures of hippocampal neurons were prepared as previously described [13] with some modifications in the protocol. Briefly, hippocampi were dissected from embryonic day 18 or 19 Sprague-Dawley rats of either sex. Tissue was then trypsinized (0.25% v/v), and mechanically dissociated in 1× HBSS (Invitrogen, Cergy Pontoise, France) containing 10mM HEPES (Invitrogen). Neurons were plated at a density of 120 × 103 cells/ml onto 18-mm diameter glass coverslips (Assistent, Winigor, Germany) pre-coated with 50 µg/ml poly-D,Lornithine (Sigma-Aldrich, Lyon, France) in plating medium composed of Minimum Essential Medium (MEM, Sigma) supplemented with horse serum (10% v/v, Invitrogen), L-glutamine (2 mM) and Na+ pyruvate (1 mM) (Invitrogen). After attachment for 3–4 h, cells were incubated in culture medium that consists of Neurobasal medium supplemented with B27 (1X), L-glutamine (2 mM), and antibiotics (penicillin 200 units/ml, streptomycin, 200 µg/ml) (Invitrogen) for up to 4 weeks at 37 °C in a 5% CO2 humidified incubator. Each week, one fifth of the culture medium volume was renewed.

### DNA constructs

The pcDNA3.1 Flag YFP hNKCC1 HA-ECL2 (NT931) was a gift from Biff Forbush (Addgene plasmid # 49063 ; http://n2t.net/addgene:49063 ; RRID:Addgene_49063; [13]. From this NKCC1-HA-Flag-mVenus plasmid, the following constructs were raised: NKCC1-HA-Δflag-ΔmVenus by truncation of the tags located on NKCC1 NTD, NKCC1-TA3-Flag-mVenus and NKCC1-TA3-ΔFlag-ΔmVenus with mutation of T203/207/212 to Alanines, and NKCC1-TA5-Flag-mVenus and NKCC1-TA5-ΔFlag-ΔmVenus with mutation of T203/207/212/217/230 to alanines. Threonine nucleotide sequence was changed to GCA for alanine substitution. The following constructs were also used: pCAG_rat KCC2-3Flag-ECL2 [13], pCAG_KCC2-3Flag-ECL2 T906/1007E [18], eGFP (Clontech), pCAG_GPHN.FingR-eGFP-CCR5TC [23] (gift from Don Arnold, Addgene plasmid # 46296 ; http://n2t.net/addgene:46296 ; RRID:Addgene_46296), homer1c-DsRed (kindly provided by D. Choquet, IIN, Bordeaux, France), WNK1 with “kinase-dead, dominant-negative domain” (WNK1-KD, D368A), “constitutively active” WNK1 (WNK1-CA, S382E) [24] (kindly provided by I. Medina, INMED, Marseille), and SuperClomeleon [25] (kindly provided by G.J. Augustine, NTU, Singapore). All constructs were sequenced by Beckman Coulter Genomics (Hope End, Takeley, U.K).

### Neuronal transfection

Neuronal transfections were carried out at DIV 13–14 using Transfectin (BioRad, Hercules, USA), according to the manufacturers’ instructions (DNA:transfectin ratio 1 µg:3 µl), with 1–2 µg of plasmid DNA per 20 mm well. Simple transfections of NKCC1-HA-Flag-mVenus plasmid concentration: 1 µg. The following ratios of plasmid DNA were used in co-transfection experiments : 1:0.4:0.4 µg for NKCC1 constructs together with GPHN.FingR-eGFP and homer1c-DsRed ; 1:0.2 µg for NKCC1 constructs with eGFP ; 0.7:0.7 µg for NKCC1 constructs with WNK1-KD or WNK1-CA, NKCC1 constructs with SuperClomeleon ; 0.7:0.7 µg KCC2 constructs with SuperClomeleon ; 0.5:0.5:0.5 µg SCLM + KCC2 + NKCC1. Experiments were performed 7– 10 days post-transfection. SPT, STORM and chloride imaging experiments were performed with Δflag-ΔmVenus NKCC1 constructs. Standard epifluorescence microscopy with Flag-mVenus NKCC1 constructs.

### Peptide treatment and pharmacology

The following peptides and drugs were used: myristoylated dynamin inhibitory peptide (50 µM; Tocris Bioscience), TTX (1 µM; Latoxan, Valence, France), R,S-MCPG (500 µM; Abcam, Cambridge, UK), S-MCPG (250 µM; HelloBio), Kynurenic acid (1 mM; Abcam), gabazine (10 µM; Abcam), muscimol (10 µM; Abcam), WNK463 (10 µM, MedChemTronica), closantel (10 µM; Sigma), bumetanide (5 µM, Abcam). R,S-MCPG and S-MCPG were prepared in equimolar concentrations of NaOH; TTX in 2% citric acid (v/v); closantel in DMSO (Sigma). Equimolar DMSO concentrations were used for control experiments in these conditions. For SPT experiments, neurons were transferred to a recording chamber, pre-incubated in presence of drugs and/or peptide at 31 °C for 10 min in imaging medium (see below for composition) and used within 45 min in presence of the appropriate drug for imaging. For immunofluorescence experiments, drugs were added directly to the culture medium for 30 min in a CO2 incubator set at 37 °C. The imaging medium consisted of phenol red-free minimal essential medium supplemented with glucose (33 mM; Sigma) and HEPES (20 mM), glutamine (2 mM), Na^+^-pyruvate (1 mM), and B27 (1X) from Invitrogen. The 138mM [Cl^−^] extracellular solution was composed of 2 mM CaCl_2_, 2 mM KCl, 3 mM MgCl_2_, 10 mM HEPES, 20 mM glucose, 126 mM NaCl, 15 mM Na^+^ methane sulfonate; the 0 mM [Cl^−^] extracellular solution was made of 1 mM CaSO_4_, 2 mM K+ - methane sulfonate, 2 mM MgSO_4_, 10 mM HEPES, 20 mM glucose, 144 mM Na^+^ - methane sulfonate.

### Live cell staining for single-particle imaging

Neurons were labelled as described previously. Briefly, cells were incubated for 8 min at 37 °C with primary antibodies against HA (rabbit, 1:250, Cell signaling Technology, cat #C29F4). After washes, cells were incubated for 1 min with F(ab’)2-Goat anti-Rabbit IgG (H+L) Secondary Antibody QDot emitting at 655 nm (1 nM; Invitrogen) in PBS (1 M; Invitrogen) supplemented with 10% Casein (v/v) (Sigma).

### Single-particle tracking and analysis

Cells were imaged as previously described using an Olympus IX71 inverted microscope equipped with a 60X objective (NA 1.42; Olympus) and a 120W Mercury lamp (X-Cite 120Q, Lumen Dynamics). Individual images of gephyrin-YFP and homer1c-DsRed, and QD real time recordings (integration time of 30 ms over 1200 consecutive frames) were acquired with an ImagEM EMCCD camera and MetaView software (Meta Imaging 7.7). Cells were imaged within 45 min following appropriate drugs pre-incubation. QD tracking and trajectory reconstruction were performed with homemade software (Matlab; The Mathworks, Natick, MA) as described in [26]. One to two sub-regions of dendrites were quantified per cell. In cases of QD crossing, the trajectories were discarded from analysis. Trajectories were considered synaptic when overlapping with the synaptic mask of gephyrin-mRFP or homer1c-GFP clusters, or extrasynaptic for spots four pixels (760 nm) away. The inclusion area was increased relatively to previous studies on KCC2 [13,18] due to the low numbers of NKCC1 trajectories recorded with a 380 nm distance. Expanding the radius did actually not change results for perisynaptic NKCC1 lateral diffusion, suggesting its clusters are located further away from synapses than KCC2 ones. Values of the mean square displacement (MSD) plot vs. time were calculated for each trajectory by applying the relation :

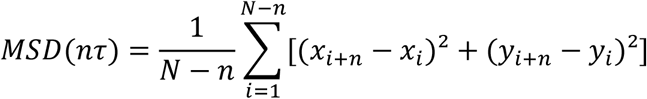

where T is the acquisition time, N is the total number of frames, n and i are positive integers with n determining the time increment. Diffusion coefficients (D) were calculated by fitting the first four points without origin of the MSD vs. time curves with the equation: MSD(nT) = 4DnT + σ ; where σ is the spot localization accuracy. Depending on the type of lamp used for imaging, the QD pointing accuracy is ∼20–30 nm, a value well below the measured explored areas (at least 1 log difference). The explored area of each trajectory was defined as the MSD value of the trajectory at two different time intervals of 0.42 and 0.45 s [27]. Synaptic dwell time was defined as the duration of detection of QDs at synapses on a recording divided by the number of exits as detailed previously [26]. Dwell times ≤ 5 frames were not retained. The number of QD vary from one cell to another and from one condition to another in a given experiment.

### Chloride imaging

Neurons were imaged at 33 °C in a temperature-controlled open chamber (BadController V; Luigs & Neumann) mounted onto an Olympus IX71 inverted microscope equipped with a 60# objective (1.42 numerical aperture (NA); Olympus). CFP and YFP were detected using Lambda DG-4 monochromator (Sutter Instruments) coupled to the microscope through an optic fiber with appropriate filters (excitation, D436/10X and HQ485/15X; dichroic, 505DCXR; emission, HQ510lp; CFP and YFP filters from Chroma Technology). Images were acquired with an ImagEM EMCCD camera (Hamamatsu Photonics) and MetaFluor software (Roper Scientific). Mean background fluorescence (measured from a nonfluorescent area) was subtracted and the ratio F480/F440 was determined. Z-stack images (16-bit; 512 × 512) were typically acquired every 30 s for 5 minutes, with an integration time of 30 ms. Regions of interest (ROIs) were selected for measurement if they could only be measured over the whole experiment.

### Immunocytochemistry

NKCC1-HA-Flag-mVenus membrane expression and clustering was assessed with staining performed after a short fixation at room temperature (RT) in paraformaldehyde (PFA; 4% w/v; Sigma) and sucrose (20% w/v; Sigma) solution in 1× PBS. The cells were then washed in PBS and incubated for 30 min at RT in goat serum (GS; 3% v/v; Invitrogen) in PBS to block non-specific staining. Neurons were then incubated for 60-180 min at RT with HA antibody (rabbit, 1:250, Cell signaling Technology, cat #C29F4) in PBS–GS blocking solution. After washing, neurons were incubated with Cy™3 AffiniPure Donkey Anti-Rabbit IgG (H+L) (1.9 µg/ml; Jackson ImmunoResearch, cat #111-165-003) for standard epifluorescence assays, or Alexa Fluor® 647 AffiniPure Donkey Anti-Rabbit IgG (H+L) (2 µg/ml, Jackson ImmunoResearch, cat #711-605-152) for super-resolution experiments, in PBS-GS solution. The coverslips were then washed, and mounted on slides. Coverslips were mounted on slides with mowiol 844 (48 mg/ml; Sigma). Sets of neurons compared for quantification were labeled and imaged simultaneously.

### Fluorescence image acquisition and analysis

Image acquisition was performed using a ×100 objective (NA 1.40) on a Leica (Nussloch, Germany) DM6000 upright epifluorescence microscope with a 12-bit cooled CCD camera (Micromax, Roper Scientific) run by MetaMorph software (Roper Scientific, Evry, France). Quantification was performed using MetaMorph software (Roper Scientific). To assess NKCC1-HA clusters, exposure time was fixed at a non-saturating level and kept unchanged between cells and conditions. For the dendritic intensity and clustering analysis, the ROI was precisely traced around focused dendrites, and global ROI pixel intensity was measured. For the clustering images were flatten background filtered (kernel size, 3 × 3 × 2) to enhance cluster outlines, and a user defined intensity threshold was applied to select clusters and avoid their coalescence. Clusters were outlined and the corresponding regions were transferred onto raw images to determine the mean NKCC1-HA cluster number, area and fluorescence intensity. For whole-dendrite intensity measurements to estimate the membrane pool fraction of NKCC1, mean pixel intensity of Venus emission and mean pixel intensity of Cy3-tagged membrane NKCC1 were computed, and then the ratio was calculated before the background flattening. The dendritic surface area of the region of interest was measured to determine the number of clusters per pixel. For each culture, we analyzed ∼10 cells per experimental condition.

### STORM microscopy

Stochastic Optical Reconstruction Microscopy (STORM) imaging on fixed samples was conducted on an inverted N-STORM Nikon Eclipse Ti microscope with a 100× oil immersion objective (NA 1.49) and an Andor iXon Ultra EMCCD camera (image pixel size, 160 nm), using specific lasers for STORM imaging of Alexa 647 (640 nm). Videos of 30,000 frames were acquired at frame rates of 50 ms. The z position was maintained during acquisition by a Nikon perfect focus system. Single-molecule localization and 2D image reconstruction was conducted as described in [28] by fitting the PSF of spatially separated fluorophores to a 2D Gaussian distribution. The position of fluorophore were corrected by the relative movement of the synaptic cluster by calculating the center of mass of the cluster throughout the acquisition using a partial reconstruction of 2000 frames with a sliding window [28]. STORM images were rendered by superimposing the coordinates of single molecule detections, which were represented with 2D Gaussian curves of unitary intensity. To correct multiple detections coming from the same Alexa 647 molecule, we identified detections occurring in the vicinity of space (2σ) and time (15 s) as belonging to the same molecule. The surface of NKCC1 clusters and the densities of NKCC1 molecules per square nanometer were measured in reconstructed 2D images through cluster segmentation based on detection densities. The minimal thresholds to determine clusters were 1% intensity, 0.1 per nm² minimum detection density and 10 detections. The resulting binary image was analyzed with the function “regionprops” of Matlab to extract the surface area of each cluster identified by this function. Density was calculated as the total number of detections in the pixels (STORM pixel size = 20 nm) belonging to a given cluster, divided by the area of the cluster.

### Statistics

Sampling corresponds to the number of quantum dots for SPT, number of cultures or animals for biochemistry, cells for ICC and chloride imaging. Sample size selection for experiments was based on published experiments, pilot studies, as well as inhouse expertise. All results were used for analysis except in few cases. For imaging experiments (chloride and calcium imaging, SPT, immunofluorescence), cells with signs of suffering (apparition of blobs, fragmented neurites) were discarded from the analysis. Data representation was usually done with boxplots or cumulative frequency plots. The statistical test to compare two groups was either Welch t-test when normality assumption was met (Q-Q plots and cumulative frequency fit), otherwise Mann-Whitney test was performed to assess the presence of a dominance or no between the two distributions. For variables following a log-normal distribution, such as variables obtained from SPT and STORM assays, we applied the log(.) function after division by the control group’s median. For super-resolution experiments, as an important variability could be observed between different cells in a same coverslip, a balanced random selection of clusters across neurons, conditions and cultures was performed, then variables from each culture were divided by the median of the control group. Results from different cultures were pooled and log(.) was applied, then the Mann-Whitney U value was computed. The process was repeated 1000 times and the p-value was determined from U distribution using the basic definition of the p-value. For SPT analysis, note that each QD is associated with 3 EA values, thus the sample size is 3 times greater. Statistical analysis were performed with R version 3.6.1 (R Core Team (2019). R: A language and environment for statistical computing. R Foundation for Statistical Computing, Vienna, Austria. Package used: ggplot2, matrixStats). Statistical tests were performed between a condition and its control only using cultures where both conditions were tested. Differences were considered significant for p-values less than 5% (*p < 0.05; **p < 0.01; ***p < 0.001; NS, not significant).

## Results

### Opposite effects of GABAergic activity on NKCC1 mobility in the axon vs. the dendrite

We questioned whether inhibitory GABAergic transmission influences NKCC1 lateral diffusion in mature (21-23 days in vitro, DIV) hippocampal cultured neurons using quantum-dot based single particle tracking (QD-SPT). We explored the impact of a pharmacological activation or blockade of GABA_A_Rs in the presence of TTX + KYN + MCPG to block glutamatergic activity and compared to the TTX + KYN + MCPG “control” condition. First, we report that in control conditions, NKCC1 diffuses along the axon (Figure S1A) and in the somato-dendritic compartment (Fig. 1A). Neurons were then acutely exposed to the GABA_A_R agonist muscimol (10 μM) or competitive antagonist gabazine (10 μM), two drugs that were shown using electrophysiological recordings to increase or block GABA_A_R-mediated inhibition in mature hippocampal cultured neurons [18]. We observed that, when exposed to gabazine or muscimol, exploration of individual QDs in the axon was increased to bigger areas, compared to QDs in control conditions (Figure S1A). The slope of the mean square displacement (MSD) as a function of time was increased for trajectories recorded in the presence of gabazine and muscimol compared to controls (Figure S1B), indicating reduced confinement of NKCC1 in the axon. This was accompanied by an increase in the diffusion coefficient (Figure S1C) and the explored area (Figure S1D). Therefore, **NKCC1 confinement is reduced in the axon under conditions of GABAergic activity blockade**.

**Fig. 1.**
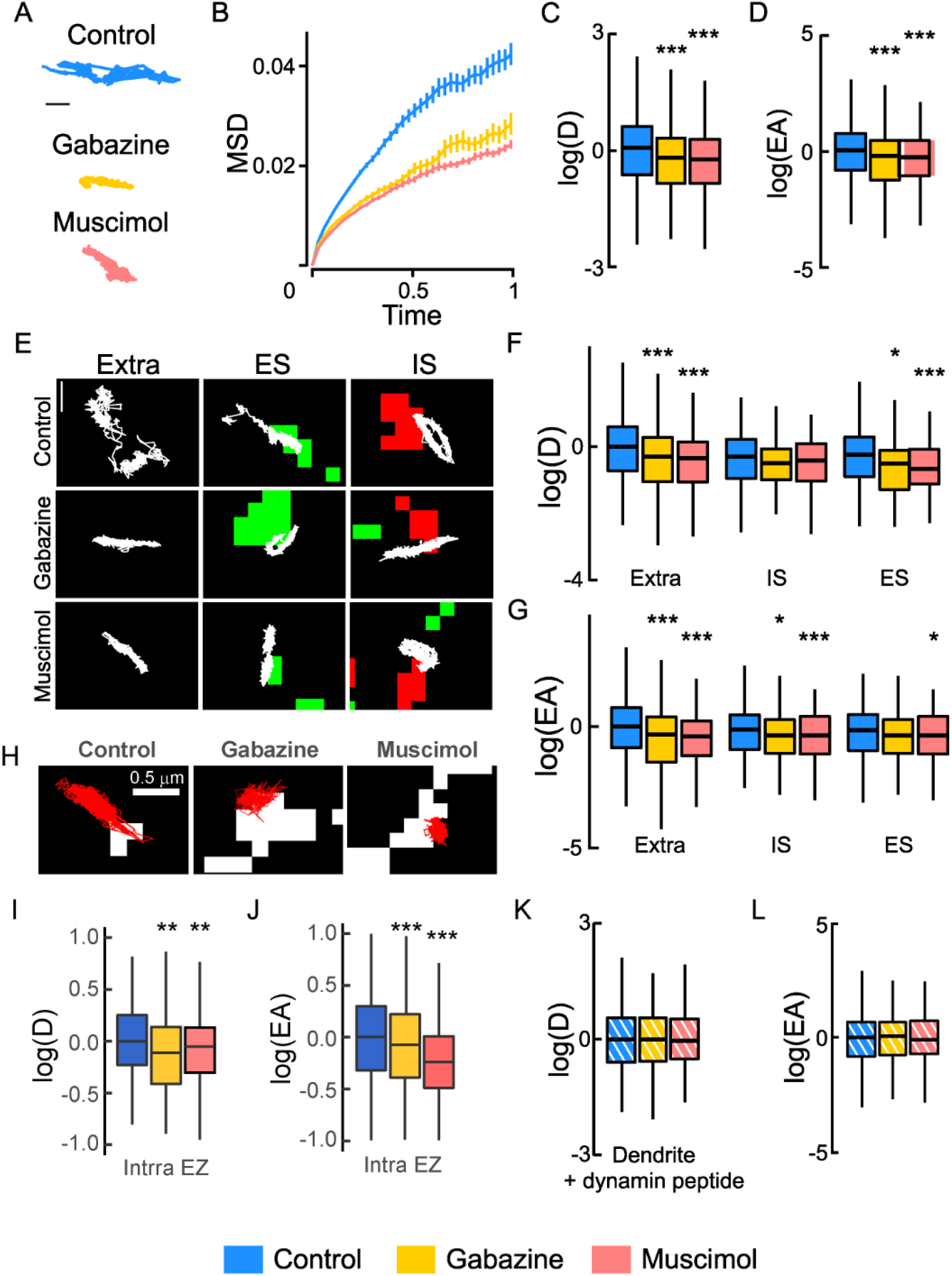
GABA_A_R activity regulates NKCC1 membrane dynamics. **A**, Examples of NKCC1 trajectories showing reduced surface exploration in the presence of gabazine or muscimol. Scale bar, 0.5 µm. **B**, Time averaged MSD functions in control (blue) vs. gabazine (yellow) or muscimol (orange) conditions show increased confinement upon gabazine or muscimol application. **C-D**, Boxplots of log(D) of NKCC1 in control condition (blue) or upon application of gabazine (yellow) or muscimol (orange) showing reduced diffusion upon gabazine or muscimol application. N = 1558 QDs (control, 41 cells), n = 387 QDs (gabazine, 18 cells), Welch t-test, p = 1.3 10^−6^, n = 545 QDs (muscimol, 27 cells), Welch t-test, p = 2.1 10^−12^, 5 cultures. **D**, Median explored area EA in control vs. gabazine or muscimol conditions show increased confinement upon gabazine (Welch t-test, p = 2 10^−8^) or muscimol (Welch t-test, p = 4.2 10^−11^) application. **E**, Trajectories (white) overlaid with fluorescent clusters of recombinant homer1c-DsRed (green) or gephyrin-Finger-YFP (red) to identify extrasynaptic trajectories (extra), trajectories at excitatory (ES) and inhibitory synapses (IS). Scale bar, 0.4 µm. **F-G**, Log(D) (**F**) and EA (**G**) of NKCC1 are decreased upon gabazine or muscimol application as compared with control condition. Note that the effect is more pronounced for extrasynaptic trajectories than for ES or IS trajectories. Diffusion coefficient (D): Extra, n = 899 QDs (control), 227 QDs (gabazine), p = 2.9 10^−5^, n = 268 QDs (muscimol) p = 2.2 10^−9^; IS, n = 244 QDs (control), 79 QDs (gabazine) p = 0.16; n = 142 QDs (muscimol) p = 0.19; ES, n = 415 QDs (control), n = 81 QDs (gabazine) p = 0.021, n = 135 QDs (muscimol) p = 0.00046. Explored area (EA): Extra, gabazine p = 2.9 10^−5^, muscimol p = 2.2 10^−9^; IS, gabazine p = 0.16, muscimol p = 0.19; ES, n = 415 QDs (control), n = 81 QDs (gabazine) p = 0.021, n = 135 QDs (muscimol) p = 0.00046. **H**, NKCC1 trajectories in control vs. gabazine or muscimol conditions in relation with endocytic zones identified by the presence of clathrin-YFP clusters. Scale bar, 0.5 µm. **I-J**, Reduced diffusion coefficient (**I**) and explored area (**J**) of NKCC1 within endocytic zones upon muscimol and gabazine treatment. Diffusion coefficient (D): Ctrl n = 117 QDs, 48 cells, Gbz n = 145 QDs, 64 cells, p = 0.0014; Ctrl n = 273 QDs, 35 cells, Musc n = 247 QDs, 28 cells, p = 0.0056, 3 cultures. Explored area (EA): Gbz, p = 5.46 10^−5^ and Musc, p = 2.2 10^−16^, 3 cultures. **K-L**, No effect of gabazine or muscimol on log(D) (**K**) and EA (**L**) of NKCC1 in conditions of blockade of endocytosis. Diffusion coefficient (D): Bulk Ctrl n = 491 QDs, Gbz n = 238 QDs, p = 0.6 and Musc n = 243 QDs, p = 0.61, 5 cultures. Explored area (EA): Gbz, p = 0.84 and Musc, p = 0.81, 5 cultures. B, F, I, K: D in µm^2^.s^-1^; C: MSD in µm^2^ vs time (s); D, G, J, L: EA in µm².

Moreover, we found that GABAergic signaling oppositely regulated the mobility of NKCC1 in the axon vs. the dendrite. In contrast to what we found in the axon, we showed that QDs explored smaller areas of the dendritic membrane following exposure to gabazine or muscimol, compared to the control condition (Fig. 1A). Analysis performed on the bulk population (extrasynaptic + synaptic) of dendritic trajectories revealed that the MSD function displayed a less steep slope for trajectories recorded in the presence of muscimol or gabazine as compared with control (Fig. 1B), indicative of increased confinement upon GABA_A_R activation or blockade. Consistent with this observation, the median diffusion coefficient and explored area values of dendritic NKCC1 were also significantly decreased upon muscimol and gabazine application (Fig. 1 C, D respectively). Thus, **the lateral diffusion of NKCC1 on the dendrite is regulated by inhibitory GABAergic transmission: the transporters being slowed down and confined in response to an increase or decrease in GABAergic activity**.

We then analyzed the effects of gabazine and muscimol on the diffusive behavior of NKCC1 in extrasynaptic and synaptic domains. Muscimol and gabazine reduced the transporter mobility and surface exploration of individual QDs (Fig. 1E). Quantitative analysis on populations of QDs revealed an impact of the treatments on diffusion coefficient and explored area both in the extrasynaptic membrane and at excitatory and inhibitory synapses (Fig. 1 F-G). The effect was greater on the diffusion coefficient than on the explored area for NKCC1 trajectories in the vicinity of excitatory synapses and vice versa for trajectories near inhibitory synapses (Fig. 1 F-G). **Thus, NKCC1 exhibits increased diffusion constraints at extrasynaptic sites and at the periphery of synapses upon GABA**_**A**_**R activation or blockade**.

### NKCC1 is targeted to endocytic zones where they are stored upon GABAergic activity changes

The increased confinement of NKCC1 induced by changes in GABA_A_R activity would result in its recruitment to membrane clusters or targeting to endocytic zones where the transporter would be stored there until later use or internalized and recycled back to the membrane or sent for degradation. We observed that acute exposure to muscimol or gabazine increased the confinement of NKCC1 in endocytic zones as observed in neurons transfected with clathrin-YFP for individual trajectories (Fig. 1H) or for hundreds of molecules (Fig. 1 I-J). Moreover, blocking clathrin-mediated endocytosis with an inhibitory peptide prevented the slow down and increased confinement of the whole NKCC1 population upon gabazine or muscimol treatment (Fig. 1 K-L). **We therefore concluded that the treatment of neurons with muscimol or gabazine increased the confinement of NKCC1 transporters in endocytic zones in the dendrites**.

To investigate whether the increased confinement of the transporter to endocytic zones induced by GABA_A_R agonists and antagonists is accompanied by an increase in its internalization, we analyzed the surface pool of NKCC1 (by calculating the ratio of the mean fluorescence intensity of the surface / surface + intracellular pool of NKCC1 (Fig. 2 A-B). The ratio remained unchanged (Fig. 2B) after exposure to muscimol or gabazine for 30 minutes, indicating that **changes in GABA**_**A**_**R activity do not affect the membrane stability of NKCC1**. This is consistent with the regulation of KCC2 lateral diffusion by GABAergic inhibition, which allows for rapid regulation of clustering and thus transporter membrane function without requiring transporter internalization [18]. We therefore examined whether changes in GABA_A_R-dependent inhibition resulted in alteration in NKCC1 clustering in hippocampal neurons. Using conventional epifluorescene, we reported that muscimol significantly reduced by 1.28-fold the density of NKCC1 clusters at the surface of transfected neurons (Fig. 2C). Furthermore, a 30 min exposure to muscimol reduced by 1.1-fold the mean size of NKCC1 clusters (Fig. 2D), as compared with untreated cells. In contrast, muscimol did not affect the mean fluorescence intensity of the clusters (Fig. 2E). These results indicate that the **increased confinement of NKCC1 in endocytic zones induced upon muscimol treatment is accompanied by a rapid reduction in its membrane clustering**. Unlike muscimol, gabazine did not noticeably alter the density of NKCC1 clusters (Fig. 2C). However, it significantly reduced the size (Fig. 2D) and intensity (Fig. 2E) of these clusters, suggesting transporter loss within clusters.

**Fig. 2.**
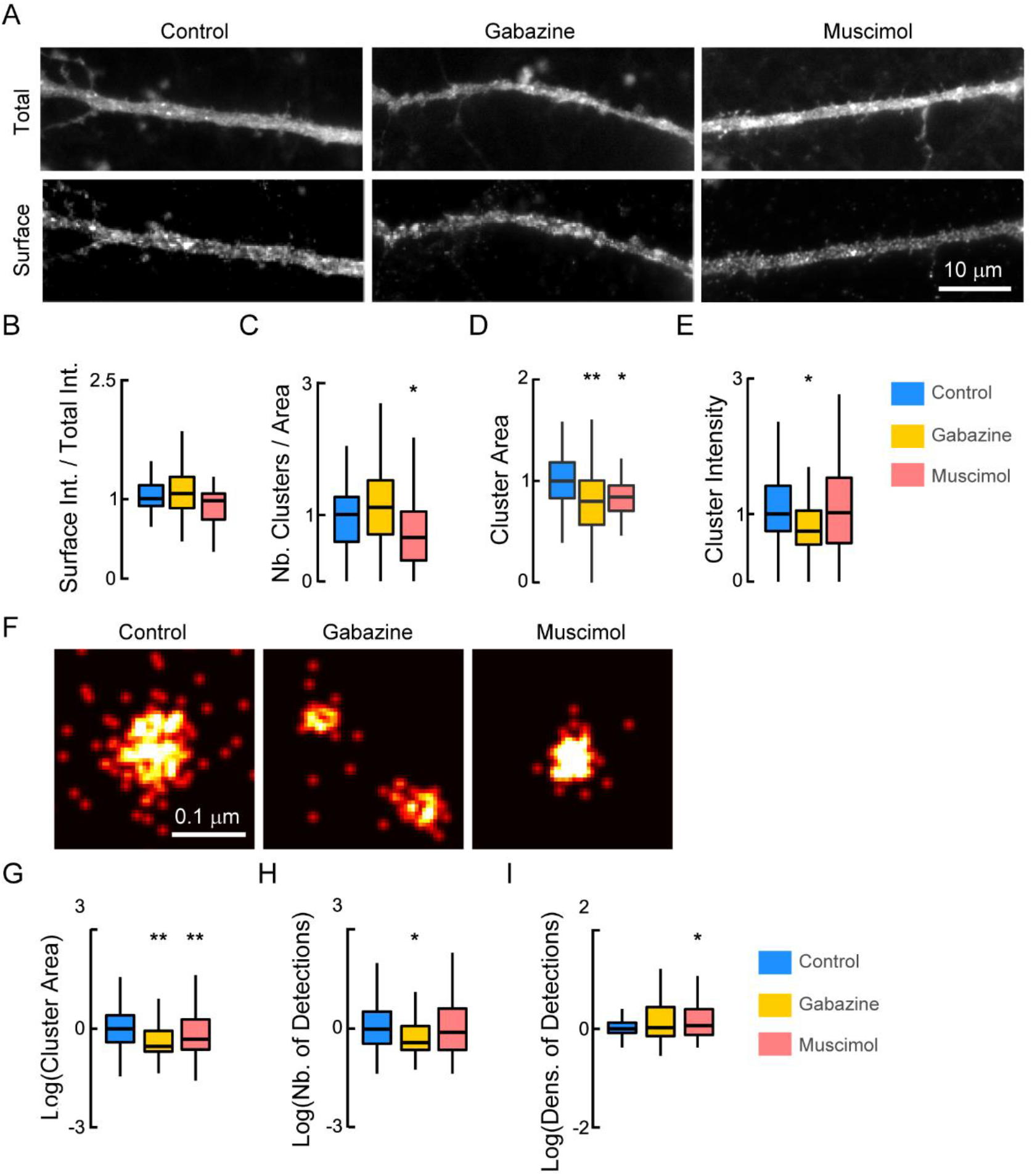
Regulation of NKCC1 membrane clustering by GABA_A_R-mediated inhibition. **A**, Conventional microscopy of HA surface staining in hippocampal neurons (DIV 21) expressing recombinant NKCC1-HA in absence (Ctrl) or presence of gabazine (Gbz), or muscimol (Musc) for 30 min. Scale bar, 10 µm. **B**, Quantification of the ratio of the surface pool of NKCC1 over the total pool of NKCC1 in control (blue), gabazine (yellow), and muscimol (orange) conditions showing no significant changes of surface NKCC1 after gabazine or muscimol treatment. Ctrl n = 30 cells, Gbz, n = 41 cells, MW test p = 0.87, Musc n = 38 cells, p = 0.17, 8 cultures. **C-E**, Quantification of NKCC1-HA cluster number (**C**), area (**D**), and intensity (**E**) shows reduced density and size of NKCC1 clusters upon muscimol treatment while gabazine treatment reduced the size and intensity of NKCC1 clusters. Values were normalized to the corresponding control values. The MW test was used for data comparison. Gabazine: cluster number (nb) p = 0.49, area p = 0.003, intensity p = 0.017. Muscimol: cluster nb p = 0.027, area p = 0.029, intensity p = 0.92. **F-I**, STORM showing that gabazine and muscimol treatments alter NKCC1 nanoclusters at the surface of hippocampal neurons. **F**, Representative STORM images of NKCC1 at the surface of neurons exposed 30 min to gabazine or muscimol. Scale bar, 0.1 µm. **G**, Quantification of NKCC1 cluster area shows reduction in nanocluster size upon gabazine and muscimol treatment. Ctrl n = 550 nanoclusters, Gbz n = 192 nanoclusters, Monte-Carlo simulations of the MW test p = 0.002, Musc n = 410 nanoclusters, p = 0.004, 4 cultures. **H**, Quantification of the number of particles detected per nanocluster showing reduced number of detection upon gabazine (Monte-Carlo simulations of MW test, p = 0.012) but not muscimol (p = 0.3) exposure. **I**, Quantification of the density of NKCC1 molecules per square micrometer highlighting denser NKCC1 packing upon neuronal exposure to muscimol (Monte-Carlo simulations of MW test, p = 0.016) but not gabazine (p = 0.97). C: µm^-1^, D: µm², G: nm², H: µm^-1^, I: µm^-2^.

Since NKCC1 cluster size is at the limit of the resolution of a standard epifluorescence microscope, we further analyzed the effect of the treatments on NKCC1 clustering using super-resolution STORM. NKCC1 form round-shaped clusters along the dendrites (Fig. 2F). We report that neuronal exposure to gabazine or muscimol altered the nanoscopic organization of NKCC1 (Fig. 2F). The muscimol treatment significantly decreased by 1.16 fold the average size of NKCC1 nanoclusters (Fig. 2G). This effect was not accompanied by a decrease in the number of molecules detected per cluster (Fig. 2H). In fact, the density of particles detected per cluster was significantly increased by 1.43 fold after muscimol exposure (Fig. 2I), indicating molecular compaction. This effect coupled with the loss of NKCC1 clusters observed with standard epifluorescence and the increased confinement of the transporter in endocytic zones **is consistent with a muscimol-induced escape of NKCC1 transporters from membrane clusters followed by their recruitment and storage in endocytic zones**.

STORM microscopy revealed that gabazine treatment induced a significant decrease in the size of NKCC1 clusters (by 0.62-fold, Fig. 2G), accompanied by a 1.5-fold reduction in the number of molecules per cluster (Fig. 2H), in agreement with the notion that transporters escaped clusters. This effect was not associated with a change in the density of molecules per cluster (Fig. 2I), reporting no change in the compaction of molecules within the cluster. **Although the effects of muscimol and gabazine differ on the nanoscale organization of NKCC1, both treatments lead to the escape of transporters from clusters, which are then rapidly captured in the endocytic zones where they are stored**.

### Intracellular chloride levels tune NKCC1 surface diffusion and clustering

We then investigated whether changes in [Cl^-^]_i_ could explain the effects of manipulations of GABA_A_R activity on the diffusion of NKCC1, as shown for KCC2 [18]. We lowered [Cl^-^]_i_ by substituting extracellular Cl^−^ with methane sulfonate in the imaging medium and tested its effect on NKCC1 diffusion. The decrease in [Cl^-^]_i_ increased NKCC1 surface exploration for individual trajectories located at the periphery of synapses and at distance (Fig. 3A). Quantification revealed that this treatment had no significant effect on the diffusion coefficient of NKCC1 on dendrites either for extrasynaptic or synaptic trajectories (Fig. 3B). On the other hand, lowering [Cl^-^]_i_ decreased the confinement of extrasynaptic transporters (Fig. 3C), while the confinement of synaptic transporters was unchanged (Fig. 3C). We then asked if the reduced confinement observed at extrasynaptic sites concerned transporters stored in endocytic zones. We found that the diffusion coefficient and surface exploration of NKCC1 were significantly increased for transporters located at distance of clathrin-coated pits (Fig. 3 D-F) while the mobility of transporters in endocytic zones was not modified upon lowering [Cl^-^]_I_ (Fig. 3 E-F). Therefore, reducing [Cl^-^]_i_ does not increase the confinement of NKCC1 in endocytic zones. **Altogether, our results provide evidence that lowering intracellular chloride levels removes diffusion constraints onto NKCC1, which move faster in the membrane, probably by being relieved from endocytic zones**.

**Fig. 3.**
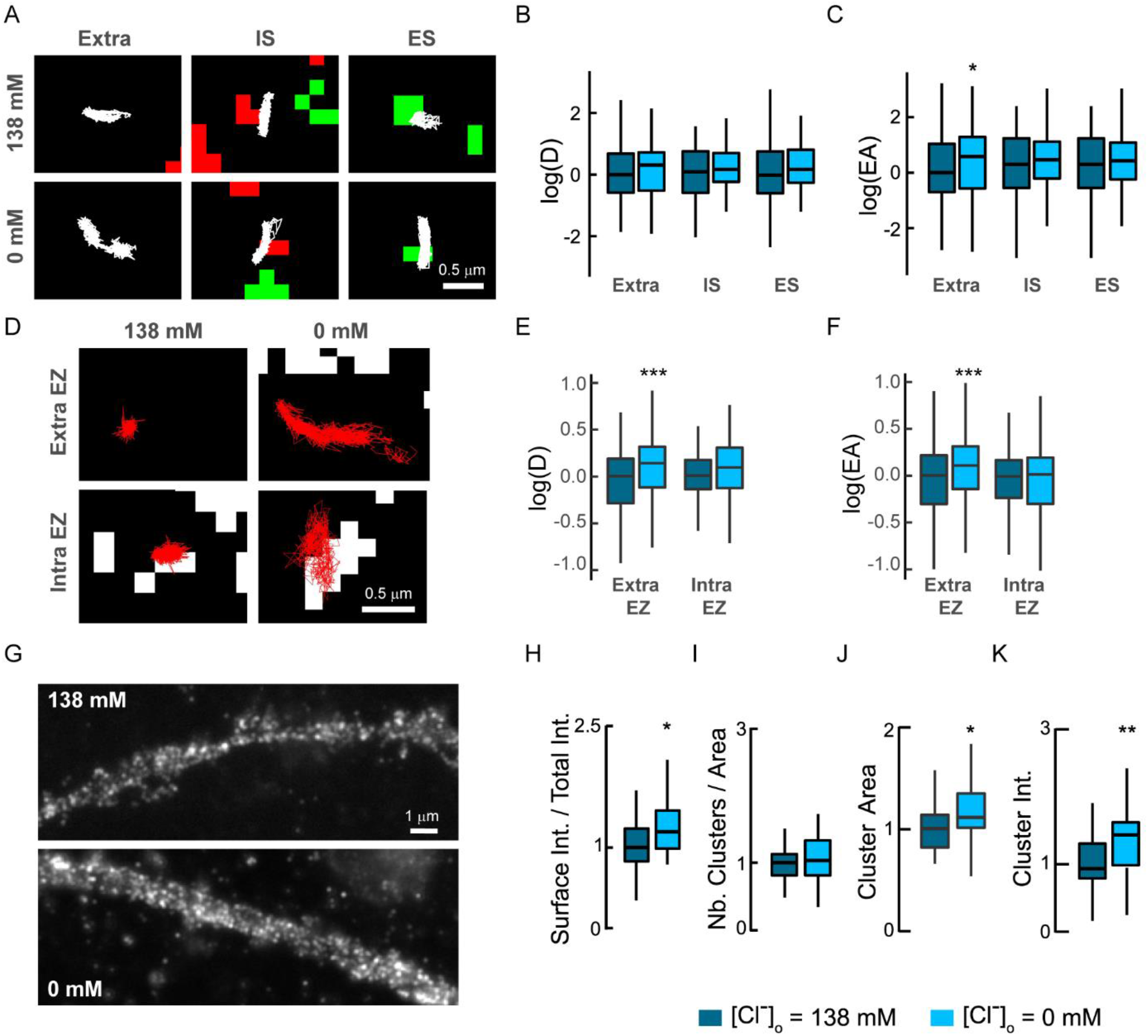
Lowering intracellular chloride increases the membrane diffusion, clustering and stability of NKCC1. **A**, NKCC1 trajectories in high vs. low extracellular Cl^-^ concentration in the extrasynaptic area (extra), and at inhibitory (IS) or at excitatory (ES) synapses. Scale bar, 0.5 µm. **B**, No major effect of a reduction of Cl^-^ concentration on log(D) (**B**) of NKCC1. Note the increase in log(EA) for extrasynaptic trajectories (**C**) in conditions of low chloride. Diffusion coefficient (D): extra, low Cl^-^ n = 128 QDs, high Cl^-^ n = 93 QDs, Welch t-test p = 0.89; IS, low Cl^-^ n = 64 QDs, high Cl^-^ n = 56 QDs, Welch t-test p = 0.74; ES, low Cl^-^ n = 62 QDs, high Cl^-^ n = 68 QDs, Welch t-test p = 0.36, 2 cultures. Explored area (EA): extra, low Cl^-^ Welch t-test p = 0.01; IS, Welch t-test p = 0.6; ES, Welch t-test p = 0.6. **D**, Examples of NKCC1 trajectories (red) inside and outside clathrin-YFP fluorescent (white) endocytic zones in conditions of low vs. high chloride extracellular Cl^-^ concentration. Scale bar, 0.5 µm. **E-F**, lowering intracellular chloride levels increases log(D) and log(EA) of NKCC1 located outside endocytic zones while the diffusion of NKCC1 inside endocytic zones is unchanged. Diffusion coefficient (D): extra EZ, low Cl^-^ n = 209 QDs, high Cl^-^ n = 238 QDs, Welch t-test p = 0.0006; intra EZ, low Cl^-^ n = 91 QDs, high Cl^-^ n = 105 QDs, Welch t-test p = 0.23, 2 cultures. Explored area (EA): extra EZ, p = 7.12 10^−8^; intra EZ, p = 0.41.**G-K**, Lowering intracellular chloride increases surface detection and clustering of NKCC1. **G**, HA surface staining in neurons expressing recombinant NKCC1-HA in in high vs. low extracellular Cl^-^ concentration for 30 min. Scale bar, 1 µm. **H**, Quantification of the ratio of the surface pool of NKCC1 over the total pool of NKCC1 in high (dark blue), and low (light blue) Cl^-^ concentration showing increase of NKCC1 surface staining upon reduction of Cl^-^ concentration. Low Cl^-^ n = 35 cells, high Cl^-^ n = 40 cells, Welch t-test p = 0.01, 3 cultures. **I-K**, Quantification of NKCC1-HA cluster number (**I**), area (**J**), and intensity (**K**) shows increased size and intensity of NKCC1 clusters upon reduction of Cl^-^ concentration. Values were normalized to the corresponding control values. Cluster Number (Nb) MW test p = 0.2, area MW test p = 0.018, intensity MW test p = 0.0056. B, E: D in µm^2^.s^-1^; C, F: EA in µm^2^ ; I : µm^-1^; J : µm².

We then determined whether this relief in diffusion constraints of the transporter was associated with its membrane redistribution. Quantification of the ratio of the surface pool of NKCC1 over the total pool of NKCC1 revealed that lowering [Cl^-^]_i_ by extracellular Cl− substitution increased NKCC1 immunoreactivity on the dendrites (Fig. 3G). This was accompanied by a 1.25 fold increase in the amount of NKCC1 detected at the cell surface (Fig. 3H). This increase in the membrane stability of NKCC1 was accompanied by an increase in its clustering. The treatment did not alter the number of clusters detected at the cell surface (Fig. 3I). However, lowering [Cl^-^]_i_ level led to a 1.12 fold increase in the median size of NKCC1 clusters (Fig. 3J) and a 1.4 fold increase in their median fluorescence intensity (Fig. 3K). These results indicate a **chloride-dependent regulation of NKCC1 diffusion-capture, consistent with a homeostatic regulation of the transporter**.

### The WNK signaling pathway regulates NKCC1

The chloride-sensitive WNK signaling pathway regulates NKCC1 activity [21]. We assessed the role of this pathway in NKCC1 clustering using overexpression of constitutively active (WNK-CA) or kinase-dead (WNK-KD) WNK1 [24] and using WNK1 (WNK463) inhibitor. The overexpression of WNK-CA had no effect on NKCC1 surface exploration and mobility for individual trajectories (Fig. 4A) and population of QDs (Fig. 4 B-C). Similarly, overexpressing WNK-CA did not alter the surface expression level of NKCC1 (Fig. 4 D-E) nor the number of NKCC1 clusters (Fig. 4F) nor the size and intensity of these clusters (Fig. 4 G-H). Based on these results, we concluded that, under basal activity conditions, **activation of the WNK1 pathway does not affect the diffusion, expression and distribution of NKCC1 at the dendritic surface in mature hippocampal neurons**.

**Fig. 4.**
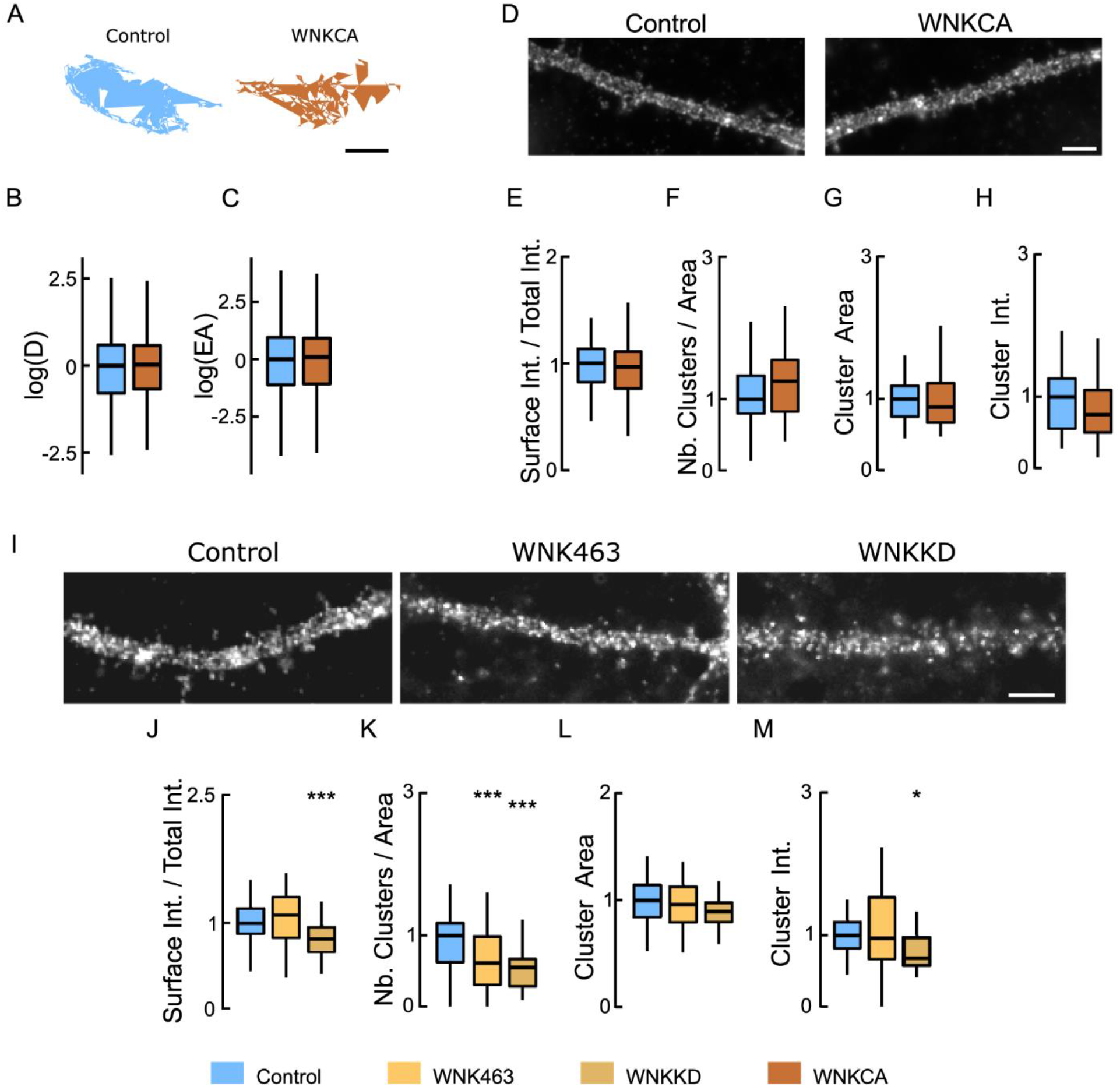
Inhibiting WNK1 reduces the clustering of NKCC1 at the neuronal surface. **A**, Representative trajectories of NKCC1 in neurons expressing constitutively-active WNK1 (WNK-CA, blue) vs. control (gray). Bar : 0.4 µm. **B-C**, WNK-CA over-expression does not change NKCC1 diffusion at the plasma membrane, with no effect on D (**B**) and EA (**C**). Diffusion coefficient (D): Bulk, Ctrl n = 305 QDs, WNK-CA n = 358 QDs, Welch t-test p = 0.81, 3 cultures; Explored area (EA): Welch t-test p = 0.49. **D**, HA surface staining in hippocampal neurons expressing recombinant NKCC1-HA together with WNK1-CA or a control plasmid. Scale bar, 4 µm. **E**, Overexpressing WNK-CA does not modify the level of NKCC1 expressed at the cell surface. Ctrl n = 45 cells, WNK-CA: n = 43 cells, Welch t-test P = 0.26, 3 cultures. **F-H**, No effect of WNK-CA expression on NKCC1 cluster number (**F**), area (**G**), and intensity (**H**). Values were normalized to the corresponding control values. Cluster Number (Nb) MW test p = 0.14, area MW test p = 0.32, intensity MW test p = 0.25. **I**, Impact of an inhibition of WNK1 on HA surface staining of neurons transfected with NKCC1-HA. WNK1 was inhibited either by incubating the neurons for 30 min with a specific inhibitor WNK-463 or by co-expressing kinase-dead WNK1 (WNK1-KD). Scale bar, 4 µm. **J**, Incubating the neurons with WNK-463 (orange) does not modify the surface level of NKCC1 compared to controls (blue), however over-expression of WNK-KD (brown) sharply reduced NKCC1 surface levels. WNK-463 experiment: Ctrl n = 24 cells, WNK-463 n = 25 cells, Welch t-test p = 0.58. WNK-KD experiment: Ctrl n = 34 cells, WNK-KD n = 24 cells, Welch t-test p = 0.0096, 4 cultures. **K-M**, Loss of NKCC1 clusters (**K**) and reduced cluster size (**L**) but not cluster intensity (**M**) upon WNK1 activity suppression with either WNK-KD or WNK-463, as compared to control conditions. Values were normalized to the corresponding control values. The MW test was used for data comparison. WNK-KD experiment: Cluster Number (Nb) p = 0.45 10^−6^, area p = 0.012, intensity p = 0.0031. WNK-463 experiment: Cluster Number (Nb) p = 0.0008, area p = 0.86, intensity p = 0.83.

Conversely, we studied the effects of an inhibition of the WNK1 signaling pathway on NKCC1 surface expression and clustering following overexpression of WNK-KD or after acute exposure to a pan-WNK antagonist (WNK-463). An acute blockade for 30 min of WNK1 with WNK463 had no effect on the membrane stability of NKCC1 (Fig. 4 I-J) while blocking WNK1 activity for 7 DIV by overexpressing WNK-KD significantly reduced the membrane stability of NKCC1 (Fig. 4 I-J). On the other hand, WNK inhibition using genetic or pharmacological approaches significantly altered the clustering of NKCC1 by decreasing respectively to 2 fold and 1.6 fold the number of NKCC1 clusters (Fig. 4K). This was not accompanied by a reduction in the size of the clusters upon WNK463 treatment or WNK-KD overexpression (Fig. 4L). However, WNK-KD overexpression decreased to 2-fold NKCC1 cluster intensity (Fig. 4M). Therefore, **inhibiting the WNK1 signaling pathway in basal activity conditions reduces NKCC1 membrane stability and clustering**.

We then studied the contribution of the WNK1 effectors SPAK and OSR1 in the regulation of NKCC1 membrane diffusion, stability and clustering using the SPAK/OSR1 inhibitor closantel [29]. An acute exposure of neurons to closantel rapidly reduced the surface explored by individual QDs (Fig. 5A). This was accompanied by a 1.14-fold reduction in NKCC1 diffusion coefficients (Fig. 5B) and by a 1.28-fold decrease in its explored area (Fig. 5C), revealing **increased NKCC1 diffusion constraints as compared with control**. This effect on diffusion was however not accompanied by a change in the surface expression of NKCC1 (Fig. 5 D-E), nor by a change in its clustering as determined by standard epifluorescence on the number of NKCC1 clusters (Fig. 5F), as well as on the size and intensity of these clusters (Fig. 5 G-H). However, the analysis of NKCC1 clusters using super-resolution imaging (Fig. 5I) revealed that a 30 min exposure to closantel reduced by 1.3-fold the cluster size (Fig. 5J). This effect was not accompanied by a significant change in the number of particles detected per cluster (Fig. 5K). However, closantel increased by 1.75-fold the density of molecules per cluster (Fig. 5L) as compared with untreated cells, indicative of molecular compaction. Therefore, the closantel-induced confinement of NKCC1 is accompanied by a rapid alteration in the nanoscale organization of the transporter. **Taken together these results show that the WNK1/SPAK/OSR1 pathway regulates the membrane dynamics, stability and clustering of NKCC1 in mature neurons. The fact that an activation of the WNK1 pathway (by overexpressing WNK-CA) has no effect on NKCC1 membrane dynamics, expression and clustering suggests that this pathway is active in mature neurons and regulates NKCC1 diffusion-capture**.

**Fig. 5.**
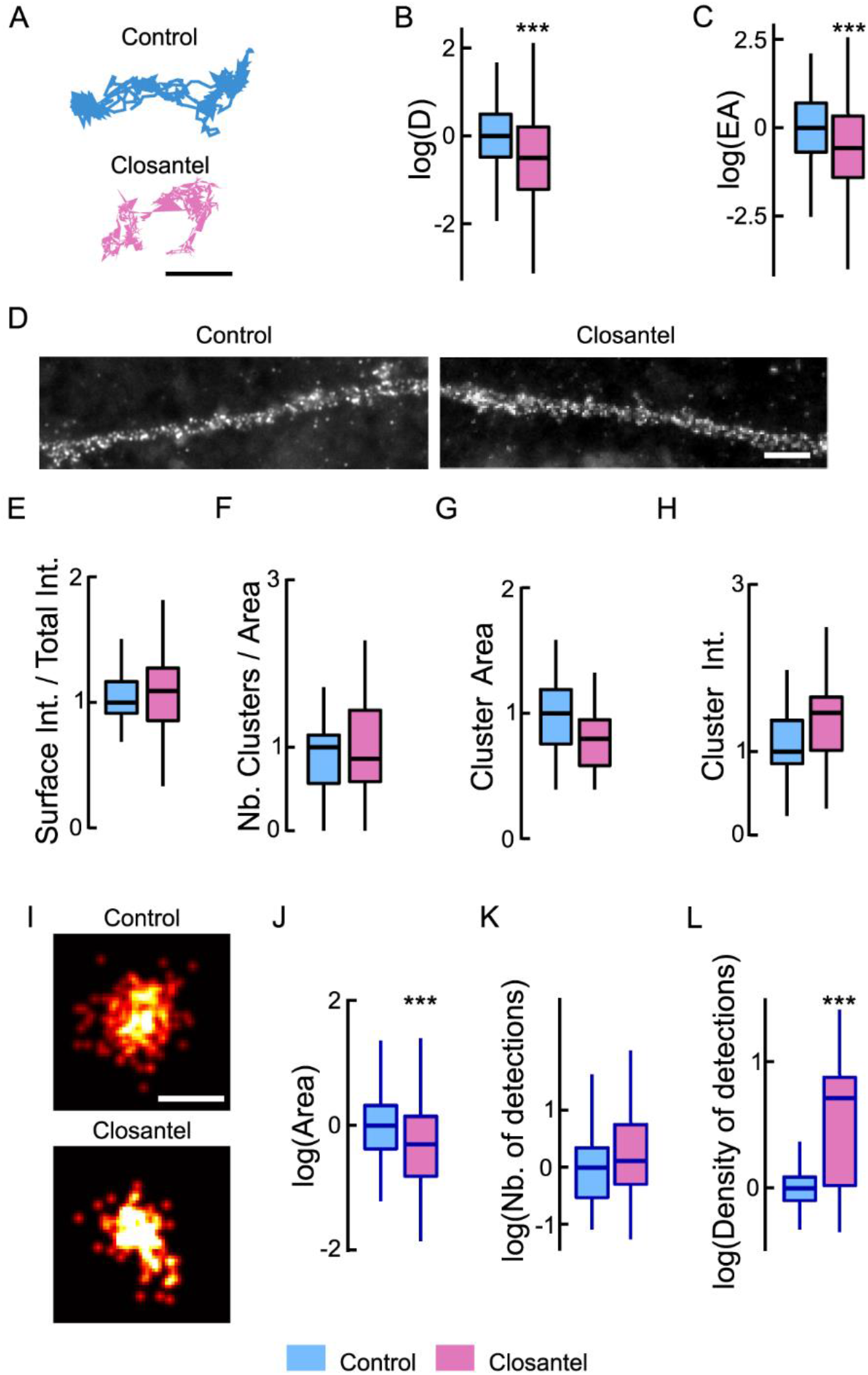
Inhibiting SPAK-OSR1 activity tunes NKCC1 membrane diffusion and clustering. **A**, NKCC1 trajectories showing reduced surface exploration in the presence of closantel. Scale bar, 0.5 µm. **B-C**, Log (D) (**B**) and EA (**C**) of NKCC1 are decreased upon closantel application (pink) as compared with control condition (blue), indicating reduced mobility and increased confinement. Diffusion coefficient (D): Bulk, Ctrl n = 433 QDs, closantel n = 371 QDs, Welch t-test p = 5.5 10^−5^, 2 cultures. Explored area (EA): Welch t-test p = 4.8 10^−11^. **D-H**, Standard epifluorescence microscopy showing no significant effect of closantel on NKCC1 membrane immunoreactivity. **D**, HA surface staining in hippocampal neurons expressing recombinant NKCC1-HA treated or not with closantel. Scale bar, 4 µm. **E**, Closantel treatment does not alter the surface expression level of NKCC1. Ctrl n = 51 cells, closantel n = 61 cells, Welch t-test p = 0.49, 7 cultures. **F-H**, Closantel has no impact on NKCC1 cluster number (**F**), area (**G**), and intensity (**H**). Cluster Number (Nb) MW test p = 0.6, area MW test p = 0.09, intensity MW test p = 0.81. **I-L**, STORM imaging showing that closantel affects NKCC1 organization at the nanoscale. **I**, Representative images of NKCC1 at the surface of neurons exposed 30 min to closantel vs control condition. Scale bar, 0.1 µm. Quantification of NKCC1 cluster area (**J**), number of particles per cluster (**K**) and density of detections in the clusters (**L**) reveal reduction in cluster size upon closantel treatment. Ctrl n = 218 clusters, closantel n = 147 clusters, 2 cultures. Cluster area: Monte-Carlo simulations of the MW test p < 0.001; Nb detection: MW test p = 0.058, density: MW test p < 0.001.

### The WNK signaling pathway targets key threonine residues on NKCC1

WNK kinases promote NKCC1 T203/T207/T212 phosphorylation [24], which in turn results in NKCC1 activation [21]. In order to test the involvement of NKCC1-T203/207/212/217/230 phosphorylation in the regulation of NKCC1 diffusion, we expressed NKCC1 constructs harboring mutations of T203/T207/T212 or T203/T207/T212/T217/T230 to alanine (TA3 and TA5, respectively) that mimic dephosphorylated states. The mobility and exploration of individual NKCC1 T203/207/212/217/230A was decreased relative to WT especially for extrasynaptic QDs (Fig. 6A). This was reflected by a 1.2 fold lower speed (Fig. 6B) and a 1.48 fold increased confinement (Fig. 6C) of QDs in the extrasynaptic membrane without changing the diffusion coefficient or the surface area explored at the inhibitory and excitatory synapses (Fig. 6 B-C). **Therefore, the dephosphorylation of NKCC1 on key threonine residues confines the transporter in the extrasynaptic membrane. In agreement with a regulation of NKCC1 by the WNK1 signaling pathway, these results indicate that a proportion of NKCC1 is phosphorylated on T203/207/212 in mature neurons. This differs from the KCC2 transporter for which regulation by the WNK1 signaling was only observed when GABA**_**A**_**R activity was challenged** [18].

**Fig. 6.**
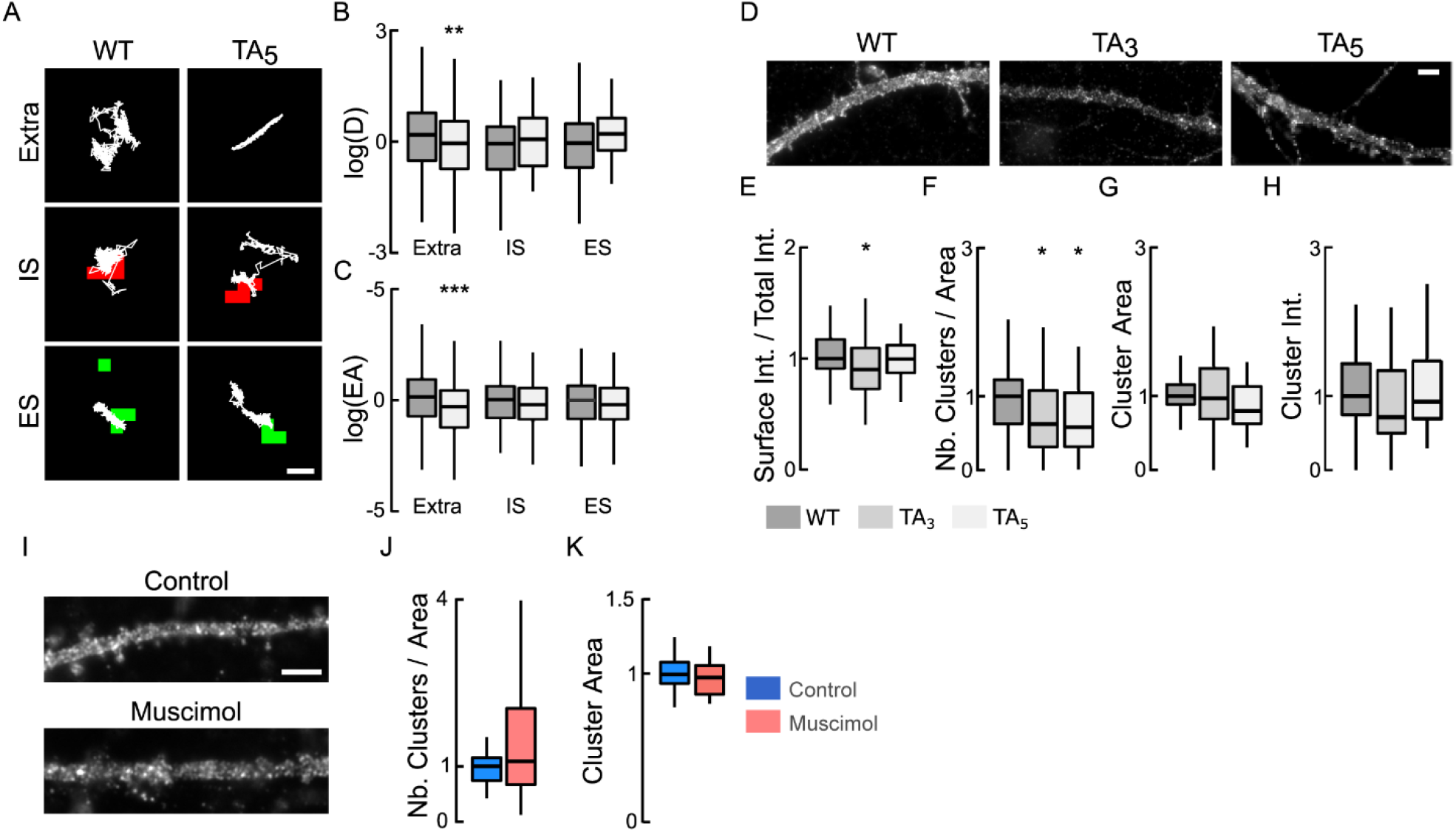
NKCC1 membrane dynamics, stability and clustering is dependent on NKCC1 phosphorylation of T203/207/212/217/230. **A**, Examples of NKCC1-T203/207/212/217/230 (WT) and NKCC1-T203/207/212/217/230A (TA_5_) trajectories in resting condition at extrasynaptic site (extra), at inhibitory (IS) and excitatory glutamatergic (ES) synapses. Scale bar, 0.5 µm. **B-C**, Log(D) (**B**) and EA (**C**) show that the dephospho-mimetic NKCC1-TA_5_ (light gray) was slower and more confined than NKCC1-WT (gray) at extrasynaptic sites but not near synapses. Diffusion coefficient (D): WT: extra, n = 189 QDs, IS, n = 42 QDs, ES n = 30 QDs; TA_5_ : extra, n = 166 QDs, IS, n = 33 QDs, ES n = 34 QDs; from 67 cells and 2 cultures. Welch t-test: extra p = 0.0088, IS p = 0.28, ES p = 0.73. Explored area (EA): extra p = 4.5 10^−4^, IS p = 0.71, ES p = 0.45. **D**, HA surface staining in hippocampal neurons expressing recombinant NKCC1-WT vs NKCC1-TA_3_ or NKCC1-TA_5_ in resting conditions. Scale bar, 4 µm. **E**, Expression of NKCC1-TA3 is slightly reduced at the plasma membrane as compared to NKCC1-WT. WT (dark gray) n = 68 cells, TA_3_ (gray) n = 43 cells, TA_5_ (light gray) n = 36 cells (9 cultures) ; WT vs TA_3_ Welch t-test p = 0.041, WT vs TA_5_ p = 0.87, 9 cultures. **F-H**, Quantification of cluster number (**F**), area (**G**) and intensity (**H**) for NKCC1-TA_3_, NKCC1-TA_5_ vs. NKCC1-WT. Note the reduced density of cluster (**F**) for NKCC1-TA_3_ and NKCC1-TA_5_ as compared to NKCC1-WT. Cluster Number (Nb) WT vs TA3 MW test p = 0.042, WT vs TA_5_ MW test p = 0.014; area: WT vs TA_3_ MW test p = 0.46, WT vs TA_5_ MW test p = 0.22; intensity WT vs TA_3_ MW test p = 0.3, WT vs TA_5_ MW test p = 0.86. **I**, HA surface staining in hippocampal neurons expressing recombinant NKCC1-TA_5_ exposed or not to muscimol. Scale bar, 4 µm. **J-K**, Muscimol application has no effect on NKCC1-TA_5_ cluster number: MW test, P = 0.45 (**J**) and area: MW test, P = 0.38 (**K**). Control: 26 cells, muscimol : 22 cells, 3 cultures.

We have previously shown in similar experimental preparations that an acute application of muscimol induces an increase in [Cl−]_i_ [18]. This is corroborated by the inhibition of WNK1 and dephosphorylation of NKCC1 [18]. Knowing that the phosphorylation of NKCC1 by WNK1 regulates its activity in non-neuronal cells [30], we wanted to know if the membrane stability and clustering of the mutated transporter was altered compared to that of the WT. Our results show that the surface pool of NKCC1 T203/T207/T212A is decreased (by 1.1 fold) compared to that of the WT transporter (Fig. 6 D-E). This decrease in the membrane stability of the transporter was accompanied by a 1.6 fold and a 1.7 fold decrease in the cluster density of NKCC1 T203/T207/T212A and NKCC1 T203/T207/T212/T217/T230A (Fig. 6F), compared to the WT. The remaining NKCC1 T203/T207/T212A and NKCC1 T203/T207/T212/T217/T230A clusters were not changed in size or fluorescence intensity compared to WT (Fig. 6 G-H). This effect is reminiscent of that observed upon muscimol treatment (Fig. 2) or WNK1 inhibition (Fig. 4). Importantly, NKCC1 T203/T207/T212/T217/T230A prevented the muscimol-induced decrease in NKCC1 clustering (Fig. 6 I-K). **We conclude that GABA**_**A**_**R-dependent regulation of NKCC1 membrane stability, diffusion and clustering involves phosphorylation of its T203/T207/T212/T217/T230 residues**.

### Functional impact of NKCC1 regulation by the WNK signaling pathway in mature hippocampal neurons

Our work describes a regulation of NKCC1 membrane stability and clustering through GABAergic activity. This regulation involves the phosphorylation of the transporter by the WNK/SPAK/OSR1 signaling pathway. To assess the functional relevance of this regulation, we looked at the impact of NKCC1 phosphorylation on [Cl^-^]_i_. We used SuperClomeleon [25] to quantify potential changes in intracellular chloride concentration. Changes of concentration were inferred from changes in YFP/CFP ratios (Fig. 7A). As a control condition, we compared the YFP/CFP ratio of cells transfected with KCC2-WT vs KCC2-T906/T1007E i.e. a construction mimicking the phosphorylated state of the transporter with a reduced capacity to extrude chloride ions, notably by modifying its membrane stability and clustering [18, 24]. An important decrease in the ratio was observed in cells transfected with KCC2-TE as compared to KCC2-WT, reflecting a higher [Cl^-^]_i_ (Fig. 7B). Thus, this approach allows the measurement of [Cl^-^]_i_ elevation resulting from manipulations of chloride co-transporter membrane expression and thereby function. We then tested whether the expression of endogenous or recombinant NKCC1 significantly impacted [Cl^-^]_i_ in our neuronal preparation. First, we determined the impact of an acute blockade of NKCC1 activity with the NKCC1 inhibitor bumetanide (5 µM) on the YFP/CFP ratio in neurons transfected with SuperClomeleon alone. The YFP/CFP ratio was comparable in both bumetanide-exposed and non-bumetanide-exposed neurons (Fig. 7C). These results are in agreement with data suggesting that **at rest, endogenous NKCC1 do not significantly influence [Cl**^**-**^]_**i**_ **in mature hippocampal neurons** [31-32]. Similarly, expression of the recombinant NKCC1-WT in mature neurons did not increase [Cl^-^]_i_ compared to neurons expressing the chloride probe alone, nor did it increase their sensitivity to bumetanide (Fig. 7C). This suggests that **the neuron tightly regulates the level of recombinant NKCC1 present at the cell membrane**.

**Fig. 7.**
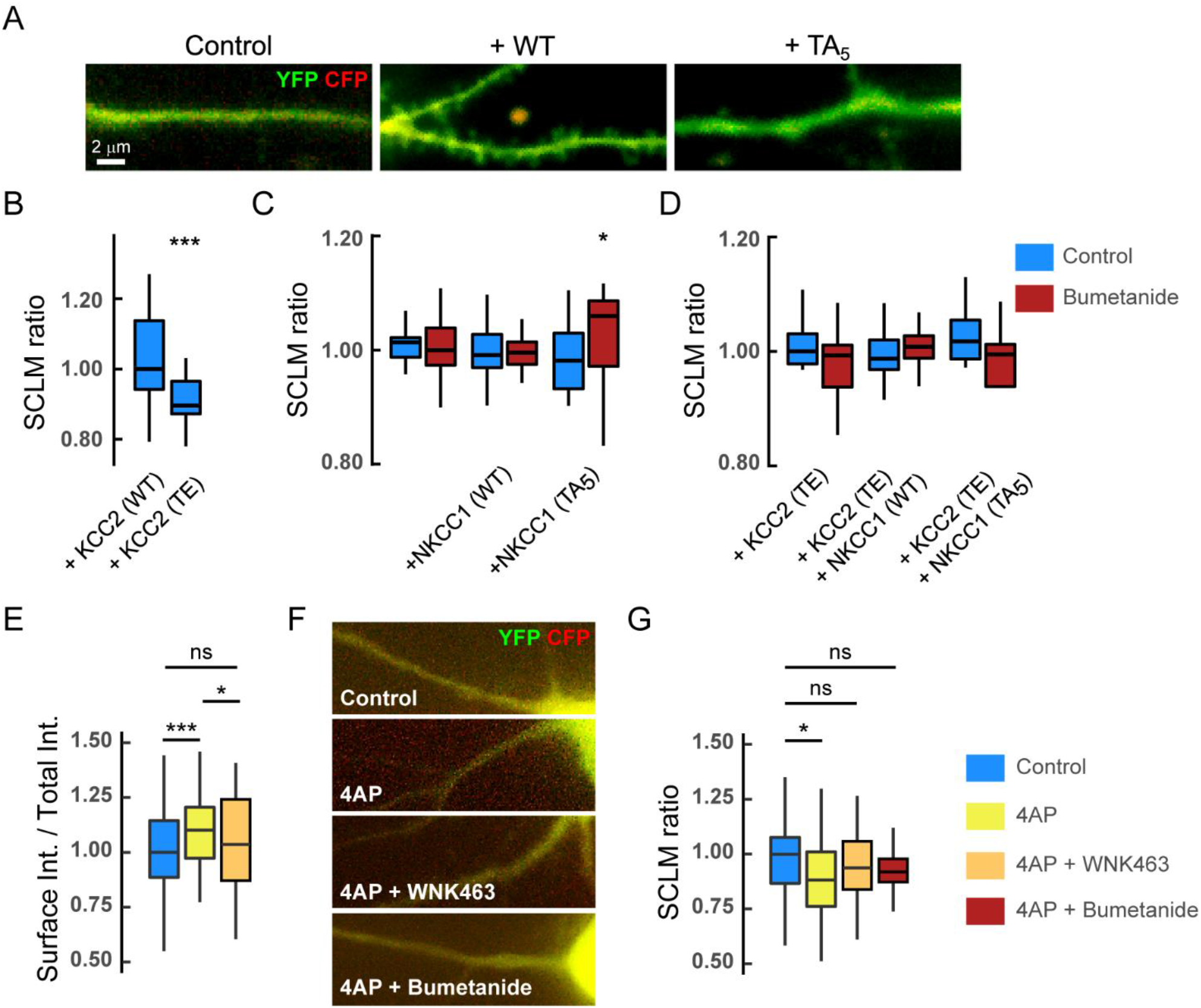
Functional impact of NKCC1 regulation by the WNK signaling on chloride homeostasis. **A-D**, In basal activity conditions, expression of recombinant WT or dephospho-mimetic NKCC1 have no major impact on [Cl^-^]_i_ in mature hippocampal neurons. **A**, composite images of CFP (red) and YFP (green) in neurons expressing SuperClomeleon (SCLM) alone (Control) or in combination with NKCC1-T203/207/212/217/230 (WT) or NKCC1-T203/207/212/217/230A (TA_5_) in resting conditions. Scale bar, 2 µm. **B**, CFP/YFP fluorescence ratio in hippocampal neurons expressing SCLM together with KCC2-T906/T1007 (WT) or phospho-mimetic KCC2-T906/T1007E (TE). KCC2-WT n = 36 cells, KCC2-TE n = 39 cells, MW test p = 7.1 10^−6^, 3 cultures. **C**, CFP/YFP fluorescence ratio in neurons expressing SCLM alone or in combination with NKCC1-WT or NKCC1-TA_5_ before (blue) and after (red) 10-30 min application of bumetanide. SCLM Ctrl n = 20 cells, Bumet n = 17 cells; SCLM + NKCC1-WT Ctrl n = 18 cells, Bumet n = 20 cells; SCLM + NKCC1-TA_5_ Ctrl n = 21 cells, Bumet n = 16 cells; 3 cultures. Ctrl vs Bumet: SCLM MW test p = 0.96, SCLM + NKCC1-WT MW test p = 0.53, SCLM + NKCC1-TA_5_ MW test p = 0.029. SCLM vs SCLM + NKCC1-WT, MW test p = 0.71, SCLM vs SCLM + NKCC1-TA_5_, MW test p = 0.69; SCLM + NKCC1-WT vs SCLM + NKCC1-TA5, MW test p = 0.89. **D**, CFP/YFP fluorescence ratio in neurons expressing the same recombinant proteins than in B but in the presence of KCC2-TE before (blue) and after (red) bumetanide treatment. SCLM + KCC2-TE Ctrl n = 15 cells, Bumet n = 15 cells, MW test p = 0.15; SCLM + KCC2-TE + NKCC1-WT Ctrl n = 13 cells, Bumet n = 16 cells, MW test p = 0.56 ; SCLM + KCC2-TE + NKCC1-TA_5_ Ctrl n = 21 cells, Bumet n = 8 cells, MW test p = 0.092; SCLM + KCC2-TE vs SCLM + KCC2-TE + NKCC1-WT, MW test p = 0.65, SCLM vs SCLM + KCC2-TE + NKCC1-TA_5_, MW test p = 0.2, SCLM + KCC2-TE + NKCC1-WT vs SuperClomeleon + KCC2-TE + NKCC1-TA_5_, MW test p = 0.12. 2 cultures. **E**, Quantification of the ratio of the surface pool of NKCC1 over the total pool of NKCC1 in control (blue), 4-AP (yellow), and 4-AP + WNK463 (orange) conditions showing a significant increase in surface NKCC1 upon 4-AP treatment, an effect prevented by pre-incubation of neurons with WNK463 (orange). Ctrl n = 63 cells, 4-AP, n = 64 cells, MW test p = 0.00046, 4-AP + WNK463 n = 57 cells, 4-AP vs. 4-AP + WNK463 p = 0.03, Ctrl vs. 4-AP + WNK463 p = 0.19, 4 cultures. **F**, Overlay images of CFP (red) and YFP (green) in neurons expressing SuperClomeleon (SCLM) and NKCC1 in control vs. 4-AP, 4-AP + WNK463 or 4-AP + Bumetanide conditions. Scale bar, 2 µm. **G**, CFP/YFP fluorescence ratio in neurons expressing SCLM and NKCC1 in control (blue) vs. 4-AP (yellow), 4-AP + WNK463 (orange) or 4-AP + Bumetanide (red) conditions. Ctrl n = 53 cells, 4-AP n = 48 cells, MW test p = 0.01; 4-AP + WNK463 n = 41 cells, MW test p = 0.20; 4-AP + Bumetanide n = 29 cells, MW test p = 0.21, 3-6 cultures.

If the expression of NKCC1-WT does not influence [Cl^-^]_i_, then it is not surprising that a loss of function of the transporter cannot be detected. Indeed, we found that NKCC1-T203/T207/T212/T217/T230A, which shows a defect in membrane clustering compared to WT, has no effect on either the YFP/CFP ratio or the response to bumetanide (Fig. 7C). However, since NKCC1 plays an important role on [Cl^-^]_i_ in mature neurons under conditions where KCC2 is down-regulated [4, 9, 18, 32], we performed additional analyses on neurons co-transfected with the mutant transporter KCC2-T906/1007E, which has a reduced chloride extrusion capacity [24] and Fig. 7A). Over-expression of KCC2-T906/1007E was preferred to the expression of a shRNA against KCC2 because this strategy led to neuronal death when NKCC1 was expressed in concert. Interestingly, neuronal death was not observed when the shRNA against KCC2 was expressed alone but only when its expression was combined with that of NKCC1 (data not shown). This indicates that the recombinant NKCC1 transporter is functional in neurons and that the influx of chloride through NKCC1 in neurons in the absence of chloride ion extrusion capacity is toxic. In agreement with previous works [31-32], this result also means that in mature neurons KCC2 is the major regulator of [Cl^-^]_i_. However, no effect of the endogenous NKCC1 or of the recombinant NKCC1-WT or NKCC1-T203/T207/T212/T217/T230A was observed on the YFP/CFP ratio, nor on bumetanide sensitivity in conditions of low KCC2 activity (Fig. 7D). We concluded **that at rest, in conditions of normal or reduced KCC2 expression, endogenous or exogenous NKCC1 transporters are not significantly contributing to [Cl**^**-**^]_**i**_ **in mature hippocampal cultured neurons**.

Since NKCC1 is overexpressed in the adult epileptic brain and that this overexpression contributes to increase seizure susceptibility [3, 7], we tested the contribution of the WNK1 signaling in the regulation of the membrane stability and function of NKCC1 in pathological conditions. For this purpose, neurons were acutely exposed to the convulsing agent 4-Amminopyridine (4-AP), a blocker of the voltage-dependent K^+^ channels responsible for membrane repolarization. This experiment was performed in absence of TTX + KYN + MCPG. The “4-AP condition” was compared to the control condition in the absence of any drug. We previously showed that an acute exposure of hippocampal neurons to 4-AP induces KCC2 endocytosis [13]. Conversely, we show here that this treatment rapidly increases the membrane expression of NKCC1 (Fig. 7E). Pre-treatment of neurons with the inhibitor WNK-463 prevented the increase in surface expression of NKCC1 induced by 4-AP (Fig. 7E), implicating WNK1 in the upregulation of NKCC1 at the neuronal surface.

Although we have shown that NKCC1 does not participate in the regulation of intracellular chloride levels under basal activity conditions in mature hippocampal neurons (Fig. 7 C-D), we show here the contribution of NKCC1 to neuronal chloride homeostasis under pathological conditions. Indeed, chloride imaging revealed that an acute exposure to 4-AP induced a significant increase in intracellular chloride levels in neurons (Fig. 7 F-G). This effect was blocked by pre-incubating the neurons with the WNK antagonist or by blocking NKCC1 activity with bumetanide (Fig. 7 F-G), thus directly implicating the WNK signaling and NKCC1 in this regulation. **Thus, we propose that 4-AP-induced hyperactivity activates the WNK pathway, which by phosphorylating NKCC1 on key threonine residues, increases its membrane expression and clustering leading to intracellular chloride influx and decreased efficacy of GABAergic transmission**.

## Discussion

We have studied, in mature hippocampal neurons, the cellular and molecular mechanisms regulating the co-transporter NKCC1, which transports chloride ions inside neurons. We showed that the transporter displays a heterogeneous distribution at the plasma membrane: it is either diffusely distributed and freely mobile in the membrane or it is organized in membrane clusters where it is slowed down and confined. Here, we show that this distribution and behavior can be rapidly tuned by GABAergic activity changes. In particular, acute GABA_A_R activation or inhibition with muscimol or gabazine respectively causes the escape of transporters from membrane clusters and its translocation to endocytic zones where it is confined. GABA_A_R-mediated regulation of NKCC1 membrane distribution uses chloride as a secondary messenger and the Cl^-^-sensitive WNK/SPAK pathway, which in turn affects the phosphorylation of a series of threonine residues on NKCC1. At rest, these modifications have little effect on [Cl^-^]_i_ but they could participate to the accumulation of Cl^-^ in neurons in pathological conditions associated with an up-regulation of NKCC1.

An increase in clustering generally correlates with a slowing down and confinement of the molecule in a sub-cellular compartment e.g. synapses for neurotransmitter receptors. Conversely, a dispersion of molecules from clusters implies a lifting of the diffusion brakes. Usually, the molecule is confined thanks to its binding to scaffolding proteins that anchors it to the cytoskeleton. However, the membrane molecule can escape from this confined region by lateral diffusion. This is the case of excitatory glutamate receptors and inhibitory GABA_A_Rs [33-34], as well as of ion transporters such as the chloride co-transporter KCC2 [11-12]. However, NKCC1 has a different diffusion behavior. The transporter was restricted in its movement when its membrane clustering was decreased (gabazine and muscimol conditions) while its diffusion was not significantly changed when its clustering increased (low chloride condition). This could be explained by the fact that a low proportion of NKCC1 transporters is clustered in the membrane while a more significant proportion of them is present in endocytic zones where they are confined and stored (in particular upon GABA_A_R activity changes).

Acute blockade of glutamatergic activity by the TTX+KYN+MCPG drug cocktail confines NKCC1 to the axon [35]. This suggests that spontaneous glutamatergic activity in contrast makes NKCC1 mobile along the axon. Here we investigated the role of GABAergic transmission on NKCC1 diffusion in the axon. We show that blocking GABA_A_R-mediated inhibition by adding gabazine to the bath in the presence of TTX+KYN+MCPG to prevent the indirect effects of gabazine on excitation, removes the constraints on NKCC1 diffusion in the axon. Conversely, activation of GABA_A_R by muscimol in the presence of TTX+KYN+MCPG slows NKCC1 in the axon. Thus, GABAergic and glutamatergic activity have opposite effects on NKCC1 diffusion in the axon. We propose that an increase in spontaneous glutamatergic activity homeostatically regulates NKCC1 diffusion in the axon to compensate for the treatment-induced increase in activity by decreasing the depolarizing/excitatory effect of GABA_A_R in the axon. Here, in contrast, muscimol-mediated GABA_A_R solicitation confines NKCC1 in the axon to enhance depolarizing GABA to counteract the increased inhibition. The effects of inhibition can also be attributed to homeostatic regulation of excitatory GABAergic transmission in the axon with potential effects on neurotransmitter release [36] and action potential firing [31].

Of note, the effects of gabazine on NKCC1 diffusion in the dendrite and axon are opposite highlighting distinct regulatory mechanisms. In the case of regulation by glutamatergic transmission, different effects were also observed in the dendrite vs. the axon. Future experiments will tell whether this difference is due to variations in intracellular chloride concentration (with a higher concentration in the axon than in the dendrite) and activation of the WNK pathway or to different molecular mechanisms.

In the dendrites of mature neurons, we observed a similar effect of GABA_A_R activation or inhibition on the diffusion and clustering of dendritic NKCC1. In both cases, the transporter was sent to endocytic zones and confined there. The fact that there was no change in the global pool (surface + intracellular) of the transporter indicates that it is stored in the endocytic zones without being internalized and degraded. These endocytic zones have been shown to constitute reserve pools of neurotransmitter receptors, which can, depending on the synaptic demand, be released and reintegrated into the diffusing pool of receptors [37]. In the case of NKCC1, this reserve pool would allow a rapid increase in the transporter availability in the plasma membrane, for example in pathological situations in which an up-regulation of NKCC1 has been observed [9].

Muscimol by activating the GABA_A_R raises [Cl^-^]_i_ [18]. An increase in [Cl^-^]_i_ inhibits the activity of WNK1 and SPAK, OSR1 kinases [38-39], leading to the dephosphorylation of NKCC1-T203/207/212/217/230 [40-42], and reduced transporter activity [43-44]. We have shown that following exposure to muscimol, NKCC1 escaped from membrane clusters and was confined in endocytic zones. Thus, transition of the transporter between membrane clusters and endocytic zones by lateral diffusion would allow modulating rapidly its availability in the membrane and its activity. The effects of muscimol are compatible with NKCC1-T203/207/212/217/230 dephosphorylation. Pharmacological (WNK-463 or closantel) or genetic (WNK-KD) blockade of the WNK/SPAK pathway or the expression of NKCC1 TA3 or NKCC1 TA5 mutants that mimic NKCC1 dephosphorylation have the same effects as muscimol: they restrict NKCC1 in their movement and reduced the membrane clustering of the transporter. The demonstration that the effect of muscimol directly involve dephosphorylation of NKCC1-T203/207/212/217/230 was provided by the fact that the effect of muscimol on NKCC1 clustering can be prevented when the mutant NKCC1-TA5 was exposed to the drug, compared to WT.

In contrast, treatment of neurons with gabazine, by blocking the activity of GABA_A_Rs, decreases dendritic [Cl^-^]_i_. In non-neuronal cells, low chloride activates WNK1/SPAK [38-39] by auto-phosphorylation of WNK1 S382 residue. Then, WNK phosphorylates in cascade SPAK on S373 and OSR1 on S325 [45], that in turn phosphorylate NKCC1 on T203/207/212/217/230 and increase the surface expression and activity of the transporter [44, 46]. We have shown that this signaling cascade is operant in mature hippocampal neurons: gabazine activates the WNK1/SPAK/OSR1 pathway by phosphorylation thus inducing the phosphorylation of KCC2-T906/1007 as well as NKCC1-T203/207/212/217/230 [18]. If muscimol decreases NKCC1 clustering and confines it to endocytic zones, gabazine treatment should conversely induce the escape of the transporter from endocytic zones thereby increasing its clustering and function in the membrane. Although we observed that a decrease in [Cl^-^]_i_ by substituting chloride with methane sulfonate decreases the confinement of NKCC1 and increases its membrane stability and clustering, treatment of neurons with gabazine did not reproduce this effect. On the contrary, gabazine confined the transporter and induced the loss of its clustering just as muscimol did. However, we have shown that antagonizing inhibition with gabazine reduces surface expression of KCC2 by increasing lateral diffusion and endocytosis of the transporter [18]. This results in reduced intracellular chloride extrusion capacity of the neuron leading to a significant increase in [Cl^-^]_i_ as monitored by SuperClomeleon imaging [18]. KCC2 is more effective in regulating [Cl-]_i_ than NKCC1 in dendrites of mature neurons [47-48]. We therefore hypothesize that the regulation of NKCC1 by gabazine is not due to a decrease in [Cl^-^]_i_ but instead to an increase in [Cl^-^]_i_ following the regulation of KCC2 by the WNK1/SPAK/OSR1 pathway. Thus, changes in the membrane expression of KCC2 (under control of the WNK1/SPAK/OSR1 pathway) would condition that of NKCC1, thus allowing [Cl^-^]_i_ to be maintained at a low level in mature neurons. This would explain why expressing recombinant NKCC1-WT in mature neurons does not significantly increase its expression at the membrane nor does it increase [Cl^-^]_i_. This suggests that the level of expression of NKCC1 at the plasma membrane is under the control of KCC2 and its tuning of [Cl^-^]_i_.

Nevertheless, we show that the membrane stability and clustering of NKCC1 can be rapidly regulated by lateral diffusion and that this mechanism is rapidly controlled by GABAergic inhibition and the WN1K/SPAK/OSR1 pathway on the dendrites of mature neurons. Although this pathway has little influence on the amount/function of NKCC1 at the neuronal surface under basal activity conditions, we propose that it may play a role in pathological situations associated with increased expression levels of NKCC1 based on our 4-AP data. Interestingly, in the pathology, upregulation of NKCC1 is often accompanied by a down-regulation of KCC2 at the neuronal surface [3, 16]. KCC2 is also regulated by diffusion-capture. We have shown that a short exposure of neurons to the convulsive agent 4-AP increases the lateral diffusion of KCC2, which escapes from the clusters, is internalized and degraded [13]. Thus, lateral diffusion would be a general mechanism to control the membrane stability of chloride co-transporters. Moreover, the fact that KCC2 is also regulated in mature neurons by the WNK1/SPAK/OSR1 pathway [18] and that this regulation has an inverse effect on membrane stability, clustering and function of KCC2 indicates that this pathway is a target of interest in the pathology. Inhibition of the pathway would prevent the loss of KCC2 and the increase of NKCC1 at the surface of the neuron, thus preventing the abnormal rise of [Cl^-^]_i_ in the pathology and the resulting adverse effects.

## Supplementary Materials

The following supporting information can be found: Figure S1: NKCC1 diffusion in the axon is increased upon GABA_A_R blockade.

## Acknowledgments

We thank J. Nabekura for kindly providing the original pEGFP–IRES–KCC2 full-length construct, D. Choquet for the homer1c–GFP construct, and P. Bregestovski for the chloride sensor CFP–YFP construct. We are grateful to the Cell and Tissue Imaging Facility of Institut du Fer à Moulin (IFM).

## Author contributions

SL conceptualized the project, designed the experiments and supervised the experimental work. SL and EC prepared the figures and wrote the paper. EC and JG performed single particle tracking experiments and analyzed the data. EC, JG and SB performed conventional wide field microscopy and analyzed the data. EC performed STORM and analyzed the data. EC and JG conducted chloride imaging and analyzed the data. MR prepared the hippocampal cultures.

## Institutional Review Board Statement

For all experiments performed on primary cultures of hippocampal neurons, animal procedures were carried out according to the European Community Council directive of 24 November 1986 (86/609/EEC), the guidelines of the French Ministry of Agriculture and the Direction Départementale de la Protection des Populations de Paris (Institut du Fer à Moulin, Animalerie des Rongeurs, license C 72-05-22).

## Informed Consent Statement

Not applicable

## Data Availability Statement

The data that support the findings of this study are available from the corresponding author upon reasonable request. The transfer of plasmids generated for this study will be made available upon request. A Materials Transfer Agreement may be required.

