## Supplementary material for "Lateral diffusion of NKCC1 contributes to neuronal chloride homeostasis and is rapidly regulated by the WNK signaling pathway": Figure S1

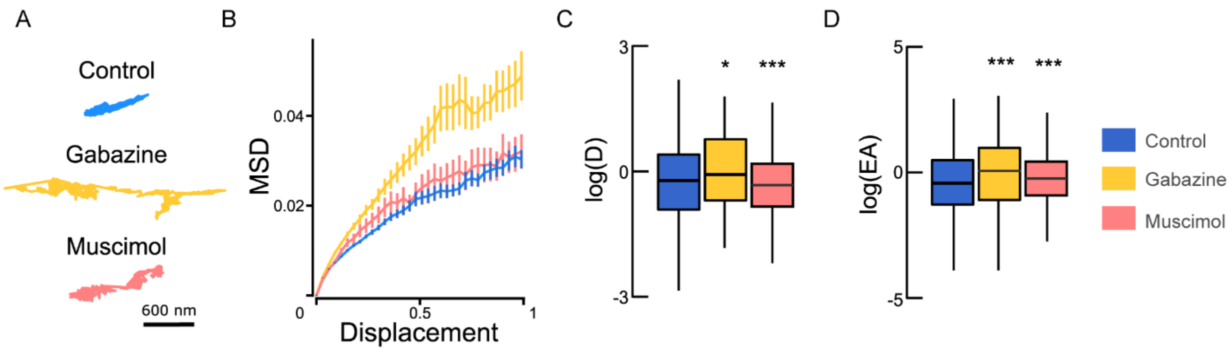


**Figure S1. NKCC1 diffusion in the axon is increased upon GABA$_A$R blockade. A,** Reconstructed trajectories of NKCC1 in the axon depicting the increased surface exploration upon gabazine (yellow) and muscimol (orange) treatment as compared to control (blue). **B,** Time-averaged MSD functions of axonal QDs in control (blue) vs. gabazine (orange) or muscimol (pink) exposure. The MSD versus time relationship for axonal trajectories shows a steeper initial slope upon drug exposure, suggesting that trajectories were less confined. $N_{Control}$ = 390 QDs, $N_{gabazine}$ = 145 QDs, $N_{muscimol}$ = 274 QDs, 37 cells from 5 cultures. **C-D**, Boxplots of log(D) of NKCC1 in control condition (blue) or upon application of gabazine (yellow) or muscimol (orange) showing increased diffusion upon gabazine application. N = 390 QDs (control, 48 cells), n = 145 QDs (gabazine, 64 cells), Welch t-test, p = 0.035, n = 274 QDs (muscimol, 28 cells), Welch t-test, p = 2.1 $10^{-12}$, 5 cultures. **D**, Median explored area EA in control vs. gabazine or muscimol conditions show reduced confinement upon gabazine (Welch t-test, p = 8.46 $10^{-6}$) or muscimol (Welch t-test, p = 4.2 $10^{-11}$) application. C: D in $\mu m^2.s^{-1}$; D: EA in $\mu m^2$.
